# Effects of long-term *in vivo* micro-CT imaging on hallmarks of osteopenia and frailty in aging mice

**DOI:** 10.1101/2020.05.10.086918

**Authors:** Ariane C. Scheuren, Gisela A. Kuhn, Ralph Müller

## Abstract

I*n vivo* micro-CT has already been used to monitor microstructural changes of bone in mice of different ages and in models of age-related diseases such as osteoporosis. However, as aging is accompanied by frailty and subsequent increased sensitivity to external stimuli such as handling and anesthesia, the extent to which longitudinal imaging can be applied in aging studies remains unclear. Consequently, the potential of monitoring individual mice during the entire aging process – from healthy to frail status – has not yet been exploited. In this study, we assessed the effects of long-term *in vivo* micro-CT imaging - consisting of 11 imaging sessions over 20 weeks - on hallmarks of aging both on a local (i.e., static and dynamic bone morphometry) and systemic (i.e., frailty index (FI) and body weight) level at various stages of the aging process. Furthermore, using a premature aging model (PolgA^(D257A/D257A)^), we assessed whether these effects differ between genotypes.

The 6^th^ caudal vertebrae of 4 groups of mice (PolgA^(D257A/D257A)^ and PolgA^(+/+)^) were monitored by *in vivo* micro-CT every 2 weeks. One group was subjected to 11 scans between weeks 20 and 40 of age, whereas the other groups were subjected to 5 scans between weeks 26-34, 32-40 and 40-46, respectively. The long-term monitoring approach showed small but significant changes in the static bone morphometric parameters compared to the other groups. However, no interaction effect between groups and genotype was found, suggesting that PolgA mutation does not render bone more or less susceptible to long-term micro-CT imaging. The differences between groups observed in the static morphometric parameters were less pronounced in the dynamic morphometric parameters. Moreover, the body weight and FI were not affected by more frequent imaging sessions. Finally, we observed that longitudinal designs including baseline measurements at young adult age are more powerful at detecting effects of *in vivo* micro-CT imaging on hallmarks of aging than cross-sectional comparisons between multiple groups of aged mice subjected to fewer imaging sessions.

## Introduction

With the estimated increase in life expectancy in the next 30 years [1], the number of people suffering from frailty will also substantially increase [2, 3]. Characterized by the decline in multiple physiological functions, which collectively result in the accumulation of health deficits, frailty leads to a higher vulnerability to adverse health outcomes including falls and osteoporotic fractures [2]. Indeed, measuring frailty, using tools such as the frailty phenotype [4] or frailty index (FI) [5], has been shown to be predictive of osteoporotic fractures [6–9]. Hence, the combined assessment of frailty and of skeletal health could be beneficial for the clinical diagnosis of osteoporosis and of frailty.

With the advent of longitudinal *in vivo* phenotyping tools such as the clinical mouse frailty index (FI) [10], animal models of frailty are of increasing interest in aging studies as the accumulation of health deficits can be quantified over time in individual animals [11–13]. Specifically, based on the frailty index in humans [14], a simple 31-item checklist including integrative measures such as grooming, strength, mobility and measures of discomfort can be used to non-invasively calculate an individualized frailty index score for each mouse [10]. Particularly owing to the striking similarities observed between the behavior of the frailty index in humans and in mice [15], the clinical mouse frailty index is considered a valuable tool to test the effects of various geroprotectors across multiple systems [13, 16, 17]. Likewise, longitudinal *in vivo* micro-CT imaging – allowing a non-invasive quantitative and qualitative assessment of the 3D bone micro-architecture over time in individual animals – has become of key importance to investigate time-dependent effects of pathologies and/or treatments in preclinical studies [18, 19]. Combined with advanced image registration techniques, bone formation as well as bone resorption activities can be directly quantified by registering consecutive time-lapsed images onto one another [20, 21]. Furthermore, the addition of computational models has been highly valuable to non-invasively estimate bone strength as well as local mechanical properties associated with aging [22, 23] and/or different interventions [24–26]. However, despite the numerous advantages of *in vivo* micro-CT imaging, the effects of cumulative radiation [18, 19], anesthesia [27–29] and handling both on a local level (i.e., on the micro-architecture and function of the scanned tissue) as well as on the systemic level (i.e., on the general well-being of the animals) must be considered. Several studies have investigated the effects of repeated *in vivo* micro-CT imaging on bone morphometric parameters in rodents, however the reports have been controversial [25, 30–36]. While *in vivo* micro-CT imaging has been shown to have dose-dependent effects on bone morphometric parameters in adolescent rats [34], studies in ovariectomized [30, 32] and adult rats [33] have shown no effects of repeated *in vivo* micro-CT imaging. Similarly, studies using mouse models have shown small but significant effects in the trabecular bone compartment [25, 30], whereas other studies reported no imaging associated effects on bone morphometric parameters [31, 35, 36]. Furthermore, the effects of repeated *in vivo* micro-CT imaging on bone morphometry have been shown to be dependent on the age of the animals, with older animals being less sensitive to imaging associated changes in bone morphometry [25]. Conversely, inhaled anesthetics have been shown to negatively affect the cognitive function in aged mice [37–39], and hence, the extent to which long-term longitudinal micro-CT imaging can be applied to monitor individual mice during the entire process of aging remains unclear.

Using an established *in vivo* micro-CT approach to monitor the 6^th^ caudal vertebrae [20, 40], we have previously shown that 5 imaging sessions did not have an effect on the bone micro-architecture and remodeling rates in 15-week old C57BL/6 mice. This approach has subsequently been used in middle-aged and aged mice [22]. By combining long-term *in vivo* micro-CT imaging of the 6^th^ caudal vertebrae with longitudinal FI measurements, we have recently identified hallmarks of frailty and senile osteoporosis in the PolgA^(D257A/D257A)^ mutator mouse [41], which, due to a defect in the proofreading activity of its mitochondrial DNA polymerase gamma, exhibits a premature aging phenotype [42, 43]. In this study, we assessed whether this long-term *in vivo* micro-CT imaging approach has biasing effects on the local and systemic level at various stages of the aging process, and whether these effects differ between genotypes. Specifically, we compared static and dynamic bone morphometric parameters as well as body weight and FI measurements of PolgA^(+/+)^ (in the following referred to as WT) and PolgA^(D257A/D257A)^ (in the following referred to as PolgA) mice subjected to 11 consecutive imaging sessions with those of mice subjected to 5 consecutive imaging sessions at various ages. By performing both cross-sectional comparisons between genotypes and between imaging groups (i.e., that were scanned at different time-points) as well as longitudinal comparisons within individual animals (i.e., that were scanned both at young adult and old age), we finally aimed to provide important insight for the effective design of studies applying *in vivo* micro-CT imaging in aging mice.

## Materials and methods

### Study design

To assess the effects of long-term *in vivo* micro-CT imaging on hallmarks of osteopenia and frailty in aging mice, we coupled longitudinal assessments of the clinical mouse frailty index with an established *in vivo* micro-CT imaging approach to monitor the changes in static and dynamic bone morphometric parameters of the sixth caudal vertebrae in a mouse model of accelerated aging, specifically the PolgA^(D257A/D257A)^ mouse. Compared to their wild-type littermates (PolgA^(+/+)^, referred to as WT), PolgA^(D257A/D257A)^ (referred to as PolgA) mice have been shown to develop multiple signs of aging by the age of 40 weeks, whereas no phenotypical differences between genotypes were observed at 20 weeks of age [41–43]. Therefore, in this study, female PolgA and WT were monitored between the ages of 20 and 46 weeks. Specifically, n=88 female mice were aged in parallel and divided into four groups (with n=12 PolgA and n=10 WT) per group. The first group was scanned between the age of 20 and 40 weeks, thus subjected to 11 bi-weekly imaging sessions over a period of 20 weeks. The other groups were subjected to our standard imaging protocol consisting of 5 bi-weekly imaging sessions over a period 8 weeks. Specifically, group 2 was imaged between weeks 26-34, group 3 between weeks 32-40 and group 4 between weeks 40-46, thus allowing continuous monitoring over the entire study period (illustrated in Fig 1). For each group, the two last scans overlapped with the two first scans of the subsequent group to allow comparison of the dynamic morphometric parameters between groups. All mouse experiments described in the present study were carried out in strict accordance with the recommendations and regulations in the Animal Welfare Ordinance (TSchV 455.1) of the Swiss Federal Food Safety and Veterinary Office and were approved by the local authorities (license numbers 262/2016, Verterinäramt des Kantons Zürich, Zurich, Switzerland).

**Fig 1.**
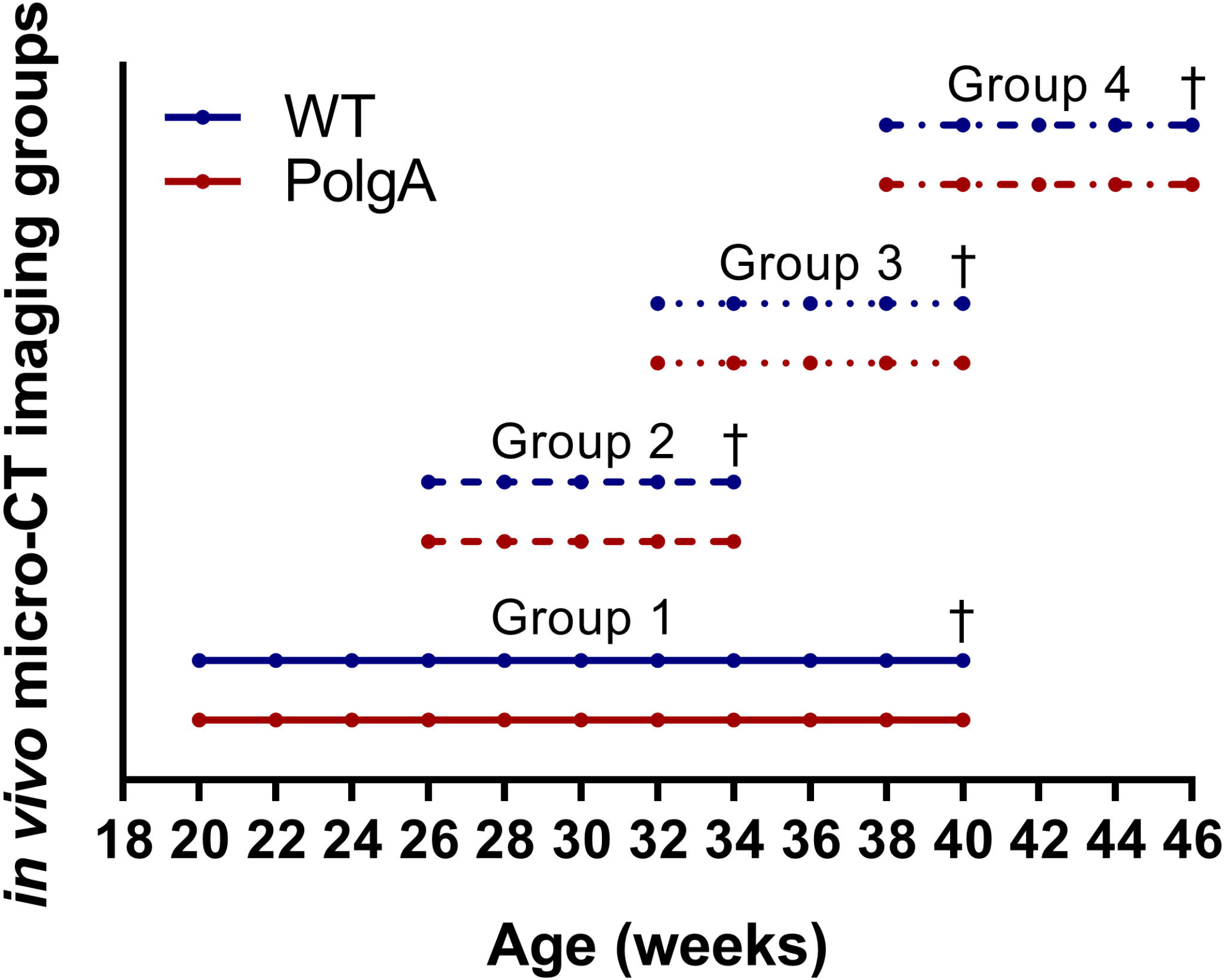
Illustration of the study design showing the time-points and duration of *in vivo* micro-CT imaging for the four groups of mice. Each group consists of n=12 PolgA (displayed in red) and n=10 WT (displayed in blue). † denotes time-point of euthanasia for the different groups, respectively.

### Animals

A colony of the mouse strain expressing an exonuclease-deficient version of the mitochondrial DNA polymerase γ (PolgA^(D257A)^, B6.129S7(Cg)-Polg^tm1Prol^/J, JAX stock 017341, The Jackson Laboratory, Farmington CT, USA) was bred and maintained at the ETH Phenomics Center (12h:12h light-dark cycle, maintenance feed and water ad libitum, 3-5 animals/cage) as previously described [41]. The presence of the PolgA knock-in mutation was confirmed by extracting DNA from ear clips (Sigma-Aldrich, KAPA Express Extract, KK7103) followed by qPCR (Bio-Rad, SsoAdvanced Universal SYBR Green Supermix, 1725272) and melt curve analysis. The primers used for genotyping (5’ to 3’; Rev Common: AGT AGT CCT GCG CCA ACA CAG; Wild type forward: GCT TTG CTT GAT CTC TGC TC; Mutant forward: ACG AAG TTA TTA GGT CCC TCG AC) were recommended by the Jackson Laboratory.

### Micro-CT imaging and analysis

*In vivo* micro-CT (vivaCT 40, Scanco Medical AG, isotropic nominal resolution: 10.5 μm; 55 kVp, 145 μA, 350 ms integration time, 500 projections per 180°, scan duration ca. 15 min, radiation dose per scan ca. 640 mGy) scans of the 6^th^ caudal vertebrae were performed every 2 weeks. The three-dimensional micro-CT images were reconstructed and registered sequentially using the algorithm described and validated by Schulte et al. [20]. After image registration, images were Gaussian filtered (sigma 1.2, support 1) and thresholded at 580 mg HA/cm^3^. Standard bone microstructural parameters were calculated in trabecular, cortical and whole bone by using automatically selected masks for these regions as described previously [40]. Dynamic morphometric parameters were calculated from the registration of consecutive micro-CT images. The voxels present only at the initial time-point were considered resorbed whereas voxels present only at the later time-point were considered formed. Voxels that were present at both time points were considered as quiescent bone. By overlaying the images, morphometrical analysis of bone formation and resorption sites within the trabecular region allowed calculations of bone formation rate (BFR), bone resorption rate (BRR), mineral apposition rate (MAR), mineral resorption rate (MRR), mineralizing surface (MS) and eroded surface (ES) [20]. Animals were anesthetized with isoflurane (induction/maintenance: 5%/1-2% isoflurane/oxygen).

### Quantification of the clinical mouse frailty index (FI)

As recommended in the recently established toolbox for the longitudinal assessment of healthspan in aging mice [16], the clinical mouse FI was quantified using the mouse frailty assessment form as published by Whitehead et al., [10], which includes the assessment of 31 non-invasive clinical items. For 29 of these items (including evaluation of the integument, the musculoskeletal system, the vestibulocochlear/auditory systems, the ocular and nasal systems, the digestive system, the urogenital system, the respiratory system, and signs of discomfort) mice were given a score of 0 if not present, 0.5 if there was a mild deficit, and 1 for a severe deficit. In addition, the mice were weighted and the body surface temperature was measured with an infrared temperature probe (6-in-1 Multifunktionsthermometer, SFT76, Silver Crest) directed at the abdomen (average of three readings was used). The later 2 items were scored based on the number of standard deviations from a reference mean in young adult mice as previously described [10]. Finally, the values were summed, and the total was divided by the number of parameters measured (i.e., 31) to provide a frailty index score between 0 and 1 for each animal. For group 1 and group 3, FI measurements were taken at 34, 38 and 40 weeks of age. For group 4, FI measurements were taken at weeks 38, 40 and 44. Although an overall high inter-rater reliability has previously been shown using the clinical mouse FI [44], all FI measurements in this study were performed by the same experienced person.

### Statistical analysis

Data are represented as mean±SD. For analysis of the longitudinal micro-CT images, frailty index and body weight measurements, linear mixed model analysis was performed for each parameter using the lmerTEST package [45] in R (R Core Team (2019). R: A language and environment for statistical computing. R Foundation for Statistical Computing, Vienna, Austria). Fixed effects were allocated to age, genotype and group and a random effect was allocated to the individual mice to account for inherent variability between mice. Furthermore, an interaction effect between age and genotype as well as between genotype and groups were assessed. Post-hoc Tukey’s multiple comparisons tests between genotypes and groups were performed using the emmeans package in R (Russell Lenth (2020). emmeans: Estimated Marginal Means, aka Least-Squares Means. R package version 1.4.4). For the comparison of the bone morphometric parameters, frailty index and body weights at 40 weeks of age, values of the individual mice are shown. Effects of genotype and imaging group as well as the interaction between genotype and group were analyzed via two-way ANOVA. In cases where a significant effect of group was found, a one-way ANOVA within groups was performed separately for each genotype followed by post-hoc Tukey’s multiple comparisons test between groupsusing SPSS (IBM Corp. Released 2016. IBM SPSS Statistics for Windows, Version 24.0. Armonk, NY, USA). Power analysis was performed in G*Power (G*Power, Version 3.1.3., Düsseldorf, Germany [46]) and in R (R Core Team (2019). R: A language and environment for statistical computing. R Foundation for Statistical Computing, Vienna, Austria).

## Results

### Effect of genotype on bone morphometry and frailty

Fig 2 shows the changes in static bone morphometric parameters over time in the different imaging groups obtained by longitudinal *in vivo* micro-CT imaging. Linear mixed model analysis revealed a significant effect of genotype for bone volume fraction (BV/TV, p<0.0001), trabecular thickness (Tb.Th, p=0.0002), trabecular number (Tb.N, p=0.002), trabecular spacing (Tb.Sp, p=0.0002), cortical area fraction (Ct.Ar/Tt.Ar, p<0.0001) and cortical thickness (Ct.Th, p<0.0001). Furthermore, the age-related changes in bone morphometric parameters developed differently between genotypes with a significant interaction effect between age and genotype for BV/TV, Tb.Th, Ct.Ar/Tt.Ar and Ct.Th (p<0.0001, Fig 2). While BV/TV, Tb.Th, Ct.Ar/Tt.Ar and Ct.Th initially increased in both genotypes, the increase in these parameters ceased to continue in PolgA mice from 30-32 weeks onwards (Fig 2A-D). On average, PolgA had lower BV/TV (−10%, p<0.0001, Fig 2A), Tb.Th (−6%, p<0.0001, Fig 2B) together with lower Tb.N (−3%, p=0.003) and higher Tb.Sp (+4%, p<0.0001) compared to WT as determined by post-hoc multiple comparisons test between genotypes. In line with the reductions in trabecular bone morphometric parameters, PolgA showed lower Ct.Ar/Tt.Ar (−6%, p<0.0001, Fig 2C) and Ct.Th (−5%, p<0.0001, Fig. 2D) compared to WT.

**Fig 2.**
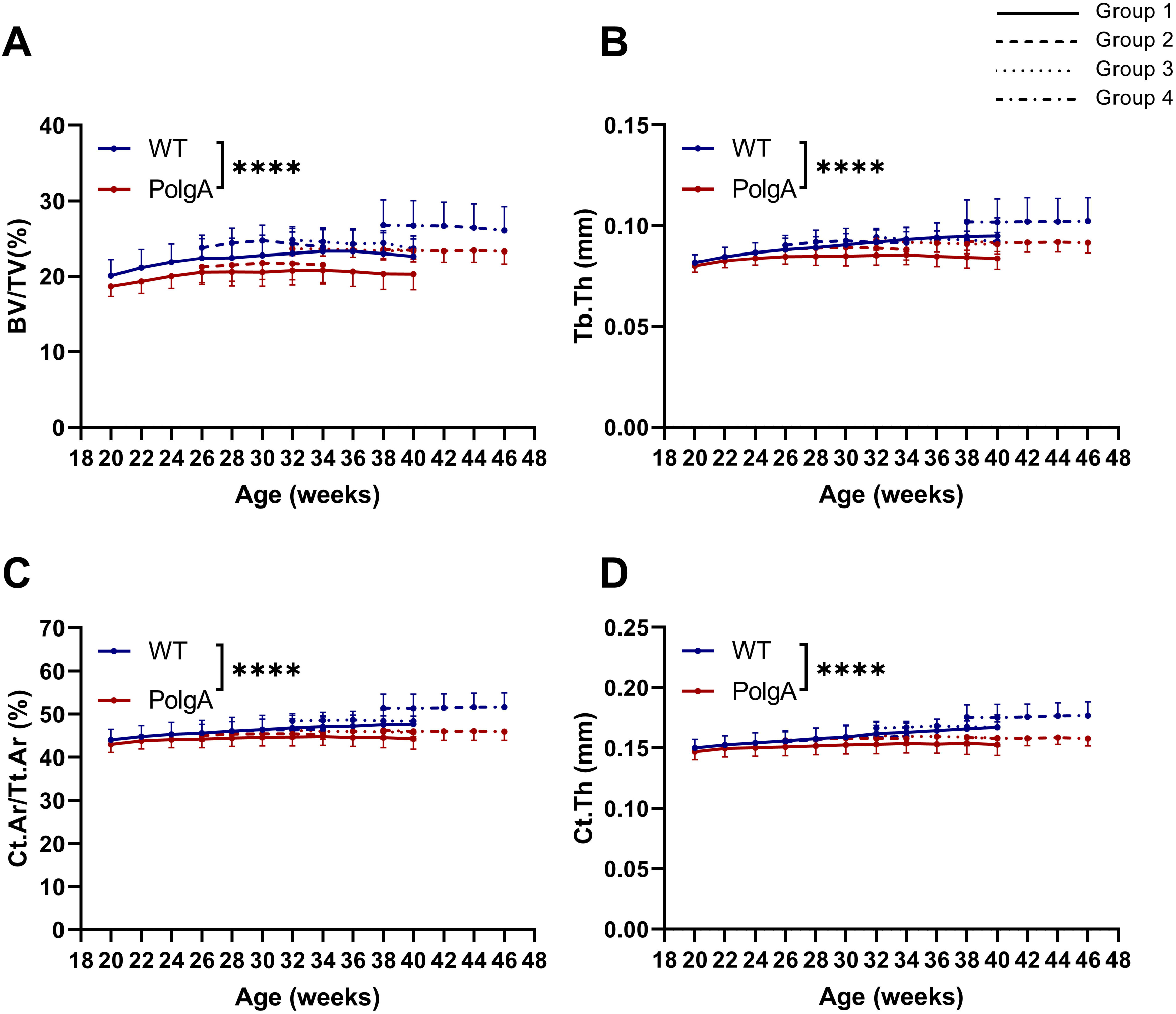
Static bone morphometric parameters obtained by longitudinal *in vivo* micro-CT monitoring of the 6^th^ caudal vertebrae between 20 and 46 weeks of age. (A) Bone volume fraction (BV/TV), (B) trabecular thickness (Tb.Th), (C) cortical area fraction (Ct.Ar/Tt.Ar) and (D) cortical thickness (Ct.Th). (**** p<0.0001 PolgA (red lines) vs WT (blue lines) determined by linear mixed model followed by post-hoc Tukey’s multiple comparisons test between genotypes. The different patterns represent different groups of mice scanned at different time-points)

By registering consecutive time-lapsed *in vivo* micro-CT images onto one-another [20], we furthermore assessed the dynamic remodeling activities in PolgA and WT mice. On average, PolgA mice had significantly lower bone formation rate (BFR, −27%, p=0.001) and bone resorption rate (BRR, −24%, p=0.001) compared to WT (Fig 3A,B). BFR did not develop differently between genotypes, whereas there was a significant interaction effect between age and genotype for BRR (p=0.014). Mineral apposition (MAR) and resorption rate (MRR), which represent the thickness of formation and resorption packages, were lower (−9%, p=0.0013 and −18%, p<0.0001) in PolgA mice compared to WT (Fig 3C,D). While MAR did not develop differently between genotypes, MRR increased in WT but remained constant in PolgA mice, thus leading to a significant interaction effect between age and genotype for MRR (p<0.001). The mineralized surface (MS), which represents the surfaces of formation sites was lower (−13%, p<0.0001) in PolgA mice compared to WT whereas ES was similar between genotypes (p>0.05, Fig 3E,F). Neither MS nor ES showed an interaction effect between age and genotype (p>0.05). Overall, these results suggest that PolgA mice have lower bone remodeling activities compared to WT.

**Fig 3.**
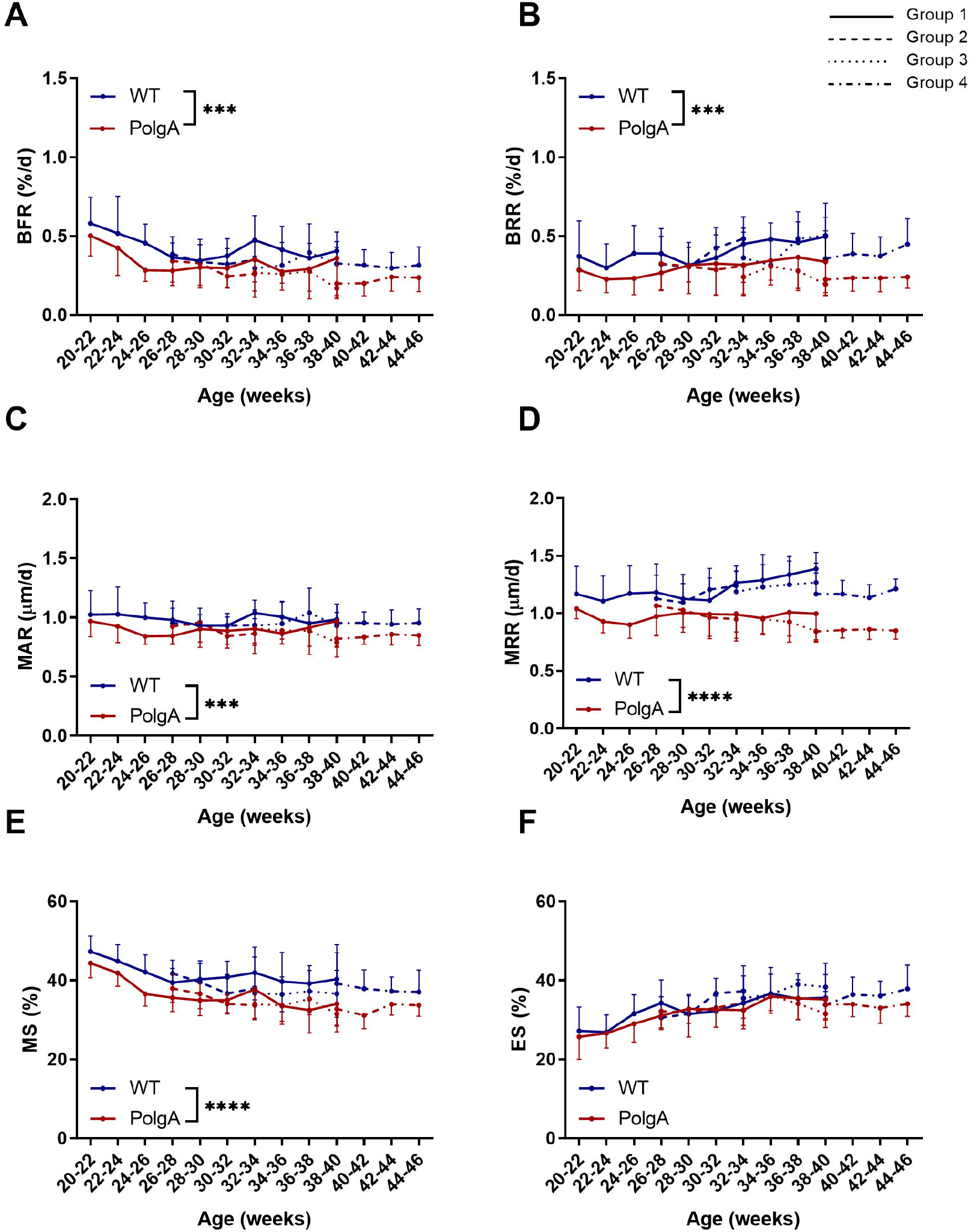
Dynamic bone morphometric parameters obtained by longitudinal *in vivo* monitoring of the 6^th^ caudal vertebrae between 20 and 46 weeks of age. (A) Bone formation rate (BFR), (B) bone resorption rate (BRR), (C) mineral apposition rate (MAR), (D) mineral resorption rate (MRR), (E) mineralizing surface (MS) and (F) eroded surface (ES). (*** p≤0.001, **** p<0.0001 PolgA (red lines) vs WT (blue lines) determined by linear mixed model followed by post-hoc Tukey’s multiple comparisons test between genotypes. The different line patterns represent different groups of mice scanned at different time-points)

Fig 4 shows the changes in clinical mouse frailty index (FI) and body weights of PolgA and WT mice in the different imaging groups. In line with the known accelerated aging phenotype of PolgA mice [42, 43], longitudinal assessments of the FI showed that the mean FI averaged over all time-points was significantly higher in PolgA (+98%, p<0.0001) compared to WT (Fig 4A). Furthermore, the FI developed differently with age in PolgA mice compared to WT (interaction effect between age and genotype, p<0.0001, Fig 4A). While PolgA and WT mice had similar FI scores at 34 weeks, PolgA mice continuously accumulated health deficits (i.e., graying, ruffled fur, distended abdomen) with age leading to higher FI scores compared to WT from 38 weeks onwards. Conversely, the body weight continuously increased in both genotypes, with no differences detected between genotypes (p>0.05, Fig 4B).

**Fig 4.**
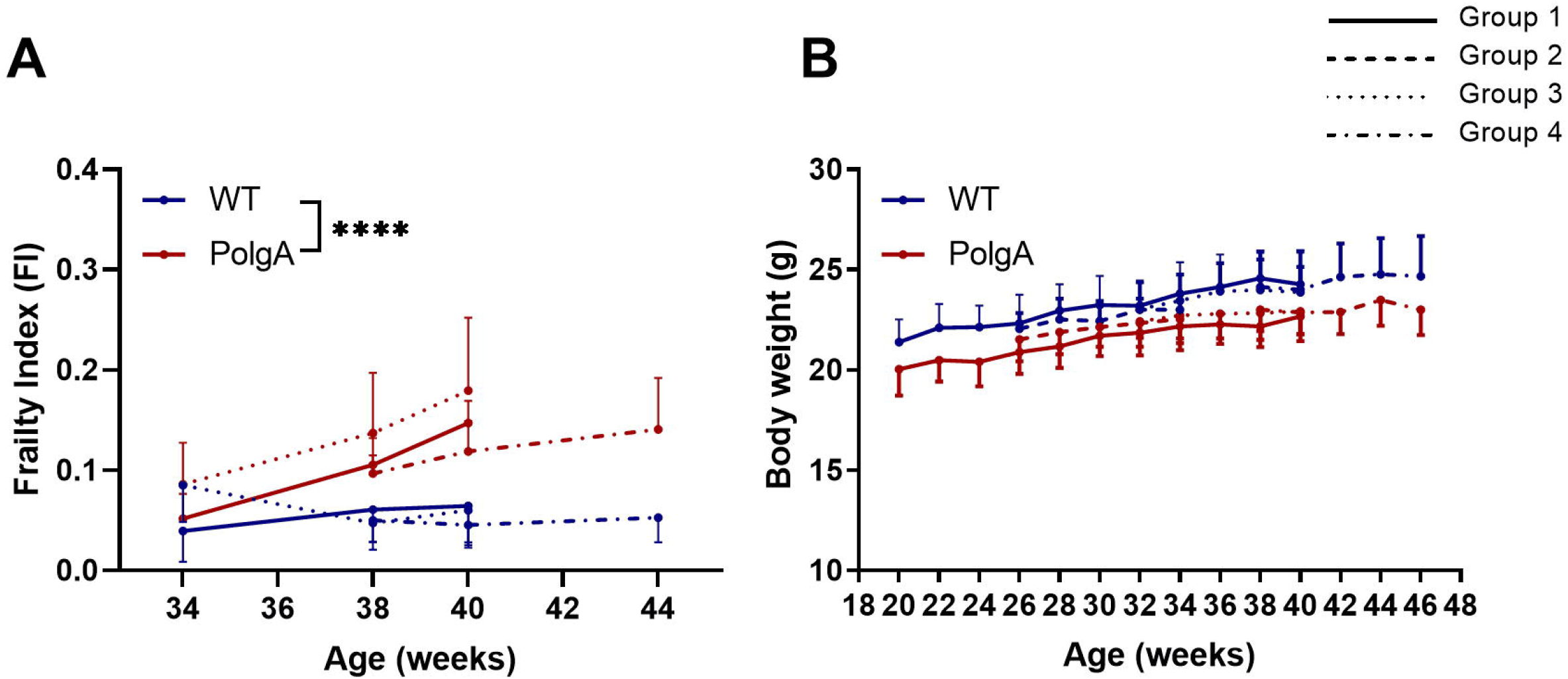
Longitudinal quantification of the (A) frailty index (FI) and (B) body weight in PolgA and WT mice between 20 and 46 weeks of age. (**** p<0.0001 PolgA (red lines) vs WT (blue lines) determined by linear mixed model followed by post-hoc Tukey’s multiple comparisons test between genotypes. The different line patterns represent different groups of mice scanned at different time-points)

### Effect of *in vivo* micro-CT imaging on bone morphometry and frailty

In order to address whether the cumulative effects of radiation, anesthesia and handling associated with long-term *in vivo* micro-CT imaging influenced the bone phenotype of PolgA and WT mice, linear mixed model analysis was performed on the static and dynamic parameters of all four groups (see study design illustrated in Fig 1). As is already visually evident in Fig 2, there was a considerable heterogeneity in static bone morphometric parameters between the different groups (effect of group for BV/TV (p<0.0001), Tb.Th (p<0.0001), Tb.Sp (p<0.0001), Ct.Ar/Tt.Ar (p<0.0001) and Ct.Th (p=0.006)). However, apart from Tb.Sp (p=0.03), none of the morphometric parameters showed a significant interaction effect between genotype and group. Post-hoc analysis between groups showed that, compared to the first group, which was subjected to 11 imaging sessions, both WT and PolgA of the fourth group had higher BV/TV (p<0.0001 for WT, p=0.009 for PolgA, Fig 2A) and Tb.Th (p=0.046 for WT, p=0.021 for PolgA, Fig 2B). The WT mice of the fourth group furthermore had higher Tb.N (p=0.04) and lower Tb.Sp (p<0.0001) compared to WT mice in the first group. With respect to the cortical bone, WT mice of the fourth group also showed higher Ct.Ar/Tt.Ar (p=0.002, Fig 2C) compared to WT mice of the first group.

The differences between groups observed in the dynamic morphometric parameters were less pronounced than in the static morphometric parameters. A significant effect of group was only found for BFR (p=0.007), BRR (p=0.01) and MRR (p=0.01, Fig 3A,B,D). Post-hoc analysis between groups showed that WT mice of the fourth group had significantly lower BRR (p=0.029) and MRR (p=0.025) compared to WT of the first group, thus explaining the higher BV/TV and Tb.Th observed in that group. Similar to the static morphometric parameters, no significant interaction effects between genotype and age were found for any of the parameters. The linear mixed model analysis of the longitudinal measurements of FI and body weights did not show any differences between imaging groups (p>0.05, Fig 4A,B).

### Effect of imaging session number on bone morphometry and frailty

In addition to evaluating the longitudinal data, the bone morphometric parameters and FI of the different imaging groups were cross-sectionally compared at 40 weeks of age, i.e. the time-point at which the different groups had received either 11 scans, 5 scans or 2 scans, respectively (Figs 5 and 6). Both genotype and imaging group significantly affected BV/TV, Tb.Th, Ct.Ar/Tt.Ar and Ct.Th (Fig 5A-D and Table S1 for p-values and effect sizes *f*), however none of the parameters showed a significant interaction effect between genotype and group. On average, PolgA mice had lower BV/TV (−8.2%, p=0.001), Tb.Th (−7.9%, p<0.0001), Ct.Ar/Tt.Ar (−7.8%, p<0.0001) and Ct.Th (−8%, p<0.0001) compared to WT. Furthermore, for Tb.Th, Ct.Ar/Tt.Ar and Ct.Th, the effect of genotype was 1.3, 1.75 and 2.5 fold stronger than the effect of imaging session number (group) as shown by the higher effect sizes *f* determined by two-way ANOVA analysis (Table S1). Conversely, for BV/TV, the effect of imaging session number was 1.48 fold stronger compared to the effect of genotype. Post-hoc analysis within genotypes showed that WT mice of the 11-scan group had lower BV/TV (p=0.008) and Ct.Ar/Tt.Ar (p=0.032) compared to those of the 2-scan group (Fig 5A,C). Similar to WT mice, PolgA of the 11-scan group showed lower BV/TV (p<0.0001) and lower Tb.Th (p=0.005 and p=0.002) compared to the 5- and 2-scan groups, with no differences between imaging groups detected in the cortical morphometric parameters (Fig 5B-D).

**Fig 5.**
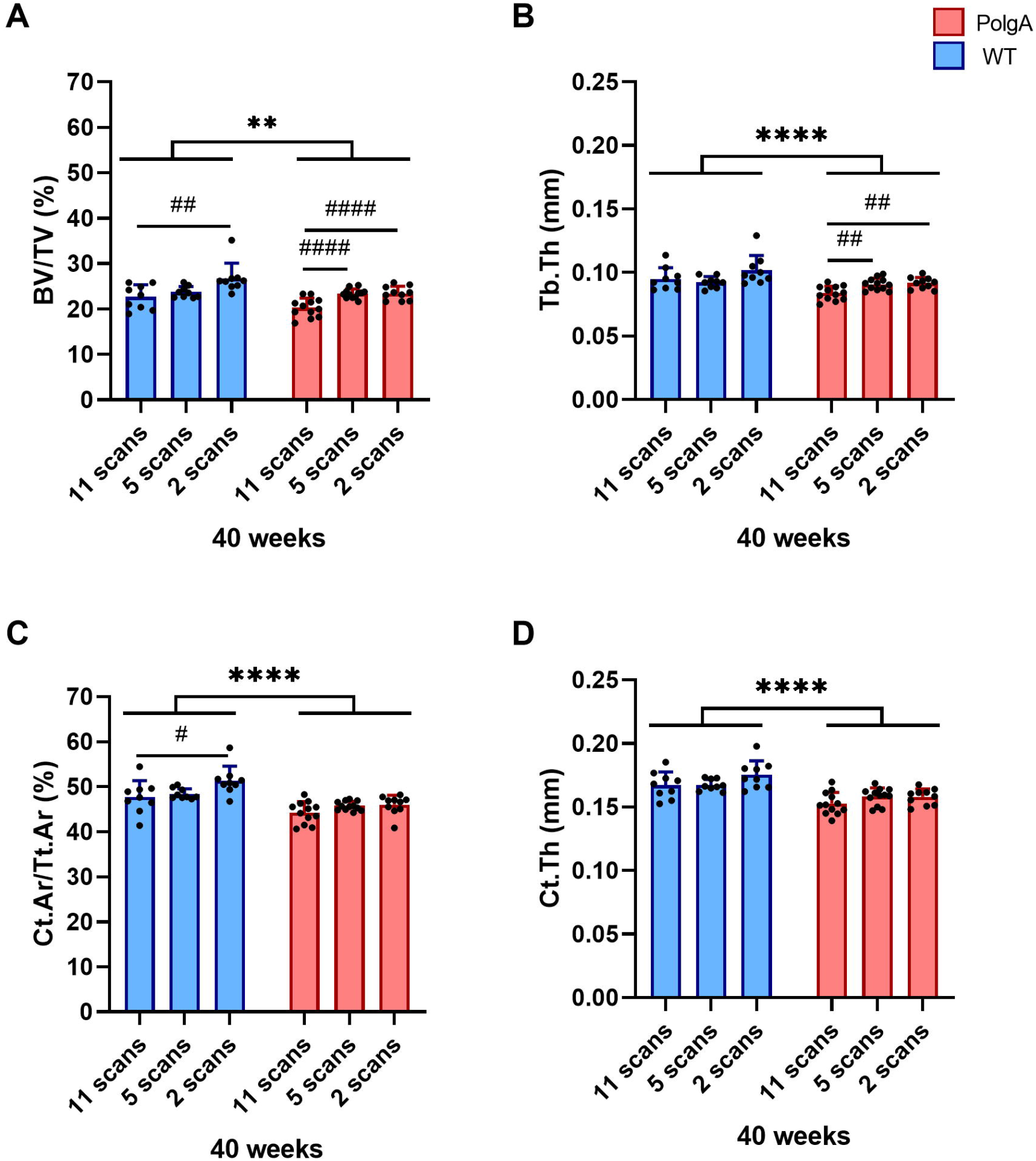
Static bone morphometric parameters at 40 weeks of age for the different groups having received either 11 scans, 5 scans or 2 scans, respectively. (A) bone volume fraction (BV/TV), (B) trabecular thickness (Tb.Th), (C) cortical area fraction (Ct.Ar/Tt.Ar), (D) cortical thickness (Ct.Th). (** p<0.01 and **** p<0.0001 between genotypes (WT (blue bars) vs PolgA (red bars)) determined by two-way ANOVA, # p<0.05, ## p<0.01 and #### p<0.0001 determined by one-way ANOVA within genotypes followed by post-hoc Tukey’s multiple comparisons test)

**Fig 6.**
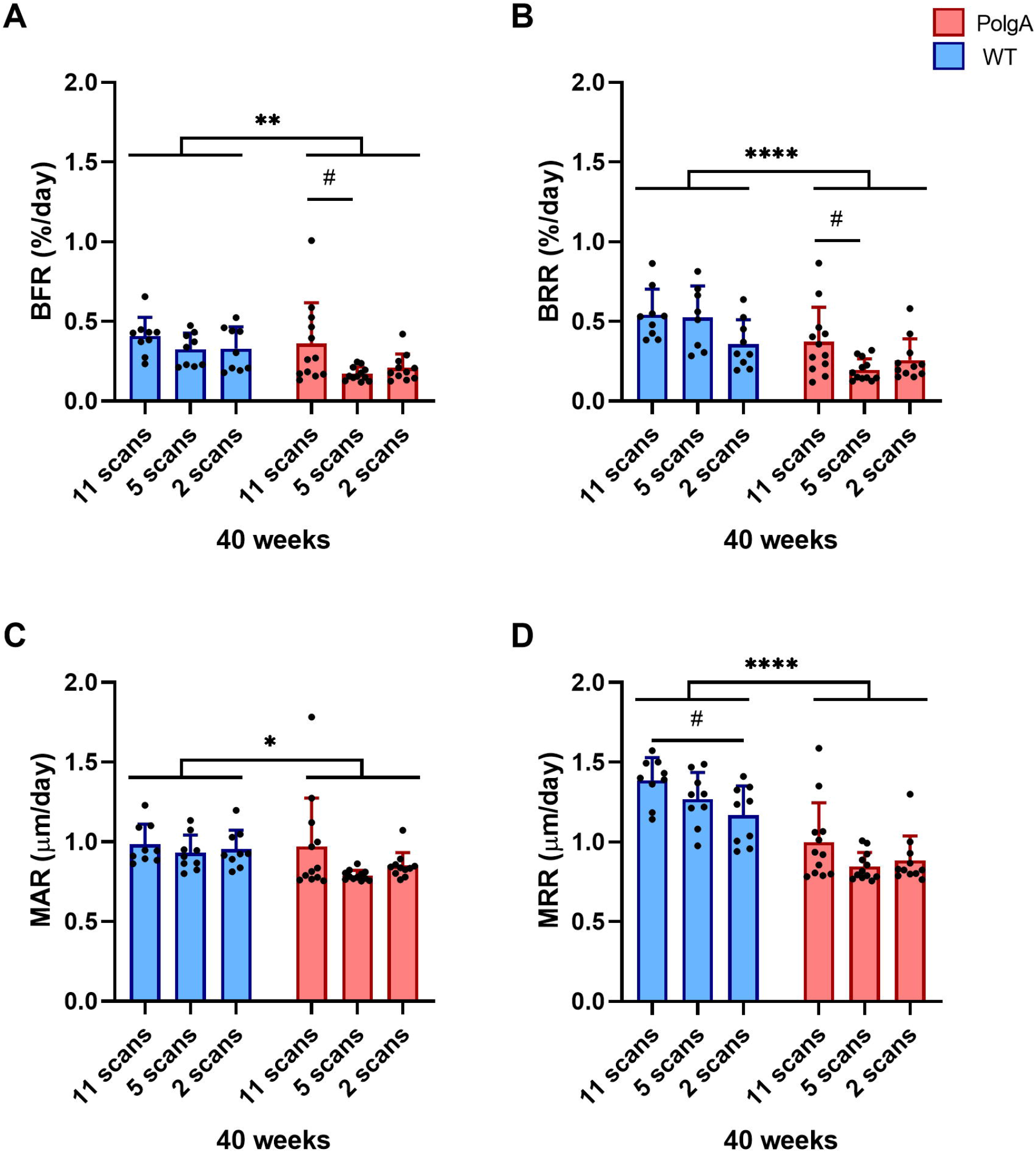
Dynamic bone morphometric parameters at 40 weeks of age for the different groups having received either 11 scans, 5 scans or 2 scans, respectively. (A) Bone formation rate (BFR), (B) bone resorption rate (BRR), (C) mineral apposition rate, (D) mineral resorption rate (MRR). (* p<0.05, ** p<0.01 and **** p<0.0001 between genotypes (WT (blue bars) vs PolgA (red bars)) determined by two-way ANOVA, # p<0.05, determined by one-way ANOVA within genotypes followed by post-hoc Tukey’s multiple comparisons test)

With respect to the dynamic parameters, both genotype and imaging session number significantly affected BFR, BRR and MRR (Fig 6A,B,D and Table S1 for p-values and effect sizes *f*), however none of the parameters showed a significant interaction effect between genotype and imaging session number. On average, PolgA mice had lower BFR (−30.8%, p=0.005), BRR (−41.9%, p<0.0001), MAR (−10%, p=0.029 and MRR (−29.7%, p<0.0001) compared to WT. Furthermore, for BRR and MRR, the effect of genotype was 1.3 and 2.4 fold stronger than the effect of imaging session number as shown by the higher effect sizes *f* determined by two-way ANOVA analysis (Table S1). Conversely, for BFR and MAR, the effect of imaging session number was 1.1 fold stronger compared to the effect of genotype. Furthermore, WT mice of the 11-scan group had higher MRR compared to WT mice of the 2-scan group (p=0.026, Fig 6H). PolgA mice of the 11-scan group had higher BFR (p=0.019) and BRR (p=0.022) compared to PolgA mice of the 5-scan group (Fig 6E,F), whereas no differences were observed between 11- and 2-scan group, respectively.

At 40 weeks of age, PolgA mice had significantly higher FI compared to WT (+166%, p<0.0001, Fig 7A). The comparison between imaging groups showed that PolgA mice in the 5-scan group had significantly higher FI scores compared to PolgA mice of the 2-scan group (p=0.019). No significant differences between genotypes or scanning groups were detected for the body weight at 40 weeks of age (Fig 7B).

**Fig 7.**
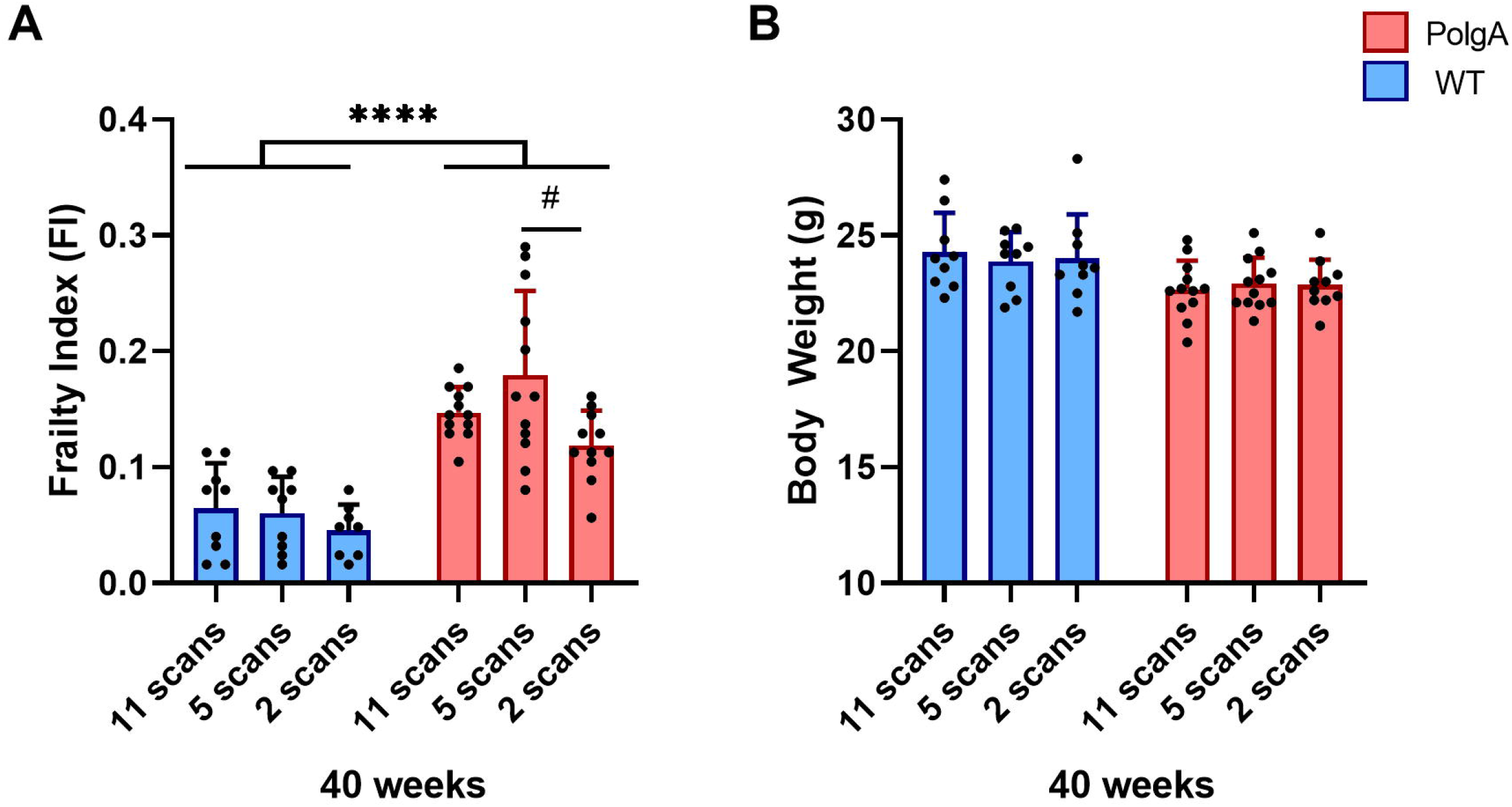
(A) Frailty index (FI) and (B) body weights at 40 weeks of age for the different groups having received either 11 scans, 5 scans or 2 scans, respectively. (**** p<0.0001 between genotypes (WT (blue bars) vs PolgA (red bars)) determined by two-way ANOVA, # p<0.05 determined by one-way ANOVA followed by post-hoc Tukey’s multiple comparisons test)

Considering the small sample sizes (n=9 per group in WT and n=12 per group in PolgA) used for this cross-sectional comparison between scanning groups, the achieved power of the analysis was computed given the obtained effect sizes (*f*) and alpha level of 0.05 (Table S2). While sufficient power (≥ 0.8) was achieved to detect any effects that might have existed in the trabecular morphometric parameters, the power for detecting differences in the cortical as well as dynamic morphometric parameters was not sufficient, suggesting that a higher number of samples would be required to detect any differences between scanning groups. Conversely, by performing a longitudinal comparison (paired t-test between parameters of individual WT and PolgA mice measured both at 20 and 40 weeks of age), sufficient power (power ≥ 0.8) was obtained for all static morphometric parameters. Nevertheless, as the dynamic bone morphometric parameters in PolgA mice remained relatively constant between 20 and 40 weeks of age, the effect sizes were small and hence, higher samples sizes would be beneficial to detect differences in dynamic bone morphometric parameters over time in individual mice.

## Discussion

Inspired by a previous study, which characterized the bone phenotype and the development of frailty in the PolgA mouse model [41], this study assessed the impact of long-term *in vivo* micro-CT imaging on hallmarks of osteopenia and frailty in individual mice during the process of aging. The unique study design, for which a large cohort of animals was aged in parallel up to 46 weeks of age, provided the possibility not only for cross-sectional comparisons between genotypes and between imaging groups (i.e., that were scanned at different time-points) but also for longitudinal comparisons within individual animals (i.e., that were scanned both at young adult and old age).

In agreement with previous studies of cross-sectional [42, 43, 47] and longitudinal designs [41], PolgA and WT had similar bone morphometric parameters at 20 weeks of age, which then diverged over time such that PolgA had significantly lower bone volume and quality at 40 weeks of age. Concomitantly, PolgA accumulated multiple health deficits over time (e.g., graying, ruffled fur, distended abdomen, kyphosis) leading to a significantly higher FI in PolgA mice from 38 weeks onwards. The clear difference in the bone morphometric parameters and FI between genotypes was observed both when groups were cross-sectionally compared and when individual mice were monitored over time. Interestingly though, *in vivo* micro-CT imaging over 20 weeks showed that this difference in bone micro-architecture was not due to bone loss in PolgA mice, but rather in the inability to reach peak bone mass, of which a more comprehensive description has been provided elsewhere [41]. Interestingly, the registration of consecutive micro-CT images revealed that PolgA mice had lower bone remodeling activities compared to WT as shown by reduced bone formation and resorption rates, with no differences in the net remodeling rate. Similarly, senile osteoporosis in humans is characterized by low bone turnover, as opposed to the high bone turnover rates (higher resorption activities) observed during postmenopausal osteoporosis [48–50].

The comparison between different imaging groups revealed a considerable heterogeneity between groups with the fourth group having higher trabecular and cortical bone morphometric parameters compared to the other groups. However, we did not observe an interaction effect between genotype and imaging groups, suggesting that the PolgA mutation does not render bone more or less susceptible to cumulative effects of radiation, anesthesia and handling associated with *in vivo* micro-CT imaging. Hence, the comparison between two genotypes remains valid despite the potential confounding effects of more frequent *in vivo* micro-CT imaging sessions. Furthermore, with the exception of BV/TV, the effect of genotype on static bone morphometric parameters was larger than the effect of repeated *in vivo* micro-CT imaging. That *in vivo* micro-CT imaging associated effects on bone morphometric parameters are stronger in trabecular bone compared to cortical bone has previously been shown [30]. In that study, the tibiae of sham- and ovariectomized-C57Bl/6J mice showed lower trabecular bone volume compared to the contralateral non-irradiated limbs, whereas this effect was not observed in healthy untreated 8- to 10-week-old control C57Bl/6J mice [30]. In line with our study, no interaction effect between ovariectomy and *in vivo* micro-CT imaging was found. This suggests that despite small but significant effects of *in vivo* micro-CT imaging on bone morphometric parameters, the comparison between different study groups/treatments remains valid. Willie et al. have also reported lower BV/TV in 10-week-old C57BL/6J mice subjected to multiple *in vivo* micro-CT scans compared to age-matched mice subjected to only one *in vivo* micro-CT scan [25]; however, this effect was not observed in 26-week old mice, suggesting that imaging associated effects are stronger in younger mice. Conversely, other studies have reported no imaging associated effects on bone morphometric parameters in mice ranging from pre-pubertal to adult age [31] up to late adulthood (48 weeks) [35]. Using the same micro-CT settings as in the present study, we have previously shown that five scans did not have an effect on the bone microstructure or bone remodeling rates in the caudal vertebrae of 15-week-old C57BL/6 mice [22, 40]. More recently, we have furthermore used a similar *in vivo* micro-CT approach to monitor specific healing phases after osteotomy and did not observe any significant imaging-associated changes in bone volume and turnover in the fracture callus of mice after 7 scans [36]. In the present study, the comparison between groups at 40 weeks of age showed that the PolgA mice that were subjected to 11 imaging sessions had lower trabecular bone morphometric parameters compared to those subjected to 5 and 2 sessions, respectively, while no differences between any of the groups were detected in the cortical bone. For the WT mice, the group subjected to 2 imaging sessions showed higher BV/TV and Ct.Ar/Tt.Ar compared to those subjected to 11 sessions, while no significant differences between 11 and 5 imaging sessions were found. Hence, although effects associated with multiple time-lapsed micro-CT scans seem to be present when compared to a very low number of 2 imaging sessions, there does not seem to be major differences between performing 5 or 11 sessions. Furthermore, although small but significant effects were observed between imaging groups in the static bone morphometric parameters, the differences between groups were less pronounced in the dynamic remodeling parameters. Taking all the longitudinal data into account, no differences between groups were observed for the parameters associated with bone formation. For parameters associated with bone resorption, the WT mice of the first group showed higher BRR and MRR compared to the fourth group. When the groups were compared at 40 weeks of age (having received either 11 or 2 imaging sessions, respectively), the 11-scan group showed higher MRR compared to the 2-scan group, suggesting a potential imaging-induced increase in osteoclast activity. An increased number and activity of osteoclasts has previously been reported in C57Bl/6 mice subjected to whole-body irradiation with an X-ray dosage of 2 Gy [51], however, as *in vivo* micro-CT imaging is localized to the scanned area, these results are not directly comparable to whole-body irradiation. One study using *in vivo* micro-CT showed dose-dependent effects on bone formation and resorption activities; while 3 consecutive scans at a high dose (776 mGy) resulted in increased bone resorption but no differences in bone formation in the tibiae of 10-week-old C57Bl/6J mice, no effects were observed when scanning at a lower dose (434 mGy) [31]. In line with these results obtained *in vivo*, scanning at the lower dose did not affect the osteogenic differentiation of bone marrow osteoprogenitors as well as osteoclast formation from bone marrow osteoclast precursors *in vitro* [31]. Although the dose used in the current study (640 mGy) lies in between the two doses reported previously, caudal vertebrae – which predominantly contain yellow (fatty) bone marrow – should be less sensitive to radiation compared to long bones, which predominantly contain red bone marrow [52, 53]. Nevertheless, owing to the low statistical power achieved in the cross-sectional comparison between imaging groups, further studies with higher animal numbers would be necessary to detect cumulative effects of long-term *in vivo* micro-CT imaging on dynamic bone morphometric parameters.

There are several limitations to this study. Firstly, we do not have a comparison to non-imaged or sham-imaged controls, nor to an internal non-irradiated control limb. However, the focus of this study was to investigate whether the long-term imaging approach comprising 11 sessions has more adverse effects compared to our standard imaging protocol comprising 5 imaging sessions, which has been shown to have no effect on the bone microstructure or bone remodeling rates in caudal vertebrae [20, 22]. In this respect, this study was designed to compare groups having been subjected to different number of imaging sessions rather than no imaging sessions at all. Nevertheless, a major limitation of this study is the fact that we do not have baseline measurements of all mice at 20 weeks of age, making it impossible to know whether the differences observed between imaging groups were due to repeated micro-CT imaging or due to initial variation between animals. In this respect, we observed that longitudinal designs including baseline measurements already at young adult age are more powerful at detecting age-related phenotypic changes compared to those including multiple groups with fewer imaging sessions. Hence, although sample sizes of 10-12 animals per group are sufficient for longitudinal studies, higher animal numbers would be beneficial for cross-sectional comparisons of aging mice. An additional limitation of this study was the inability to single out the effects of radiation, anesthesia and handling, respectively as the main factor influencing bone morphometric parameters, as more frequent imaging sessions comprise more frequent anesthesia and handling as well as more frequent fixation in the mouse holder used for imaging. Therefore, in order to rule out potential harmful effects of the imaging procedure, future studies should include a control group, receiving a baseline and end-point *in vivo* scan as well as sham-scans for the rest of the measurements. Indeed, a previous study, which included sham-scanning did not observe any imaging associated effects on the bone micro-architecture and cell viability of bone marrow cells in aged rat tibiae subjected to 8 weekly scans [33]. Furthermore, although this study assessed the effects of long-term *in vivo* micro-CT imaging on bone remodeling activities, thereby providing indirect information on cellular activities and recruitment, potential damaging effects of this longitudinal micro-CT imaging approach were not assessed at the cellular level. Future studies should therefore include analyses of gene and protein expression in response to long-term *in vivo* micro-CT imaging.

Lastly, as radiation exposure has been implicated in the degradation of mechanical properties of bone tissue [54, 55], an additional limitation of this study is the lack of end-point mechanical testing of the bone samples. A previous study, which investigated the effects of *ex vivo* ionizing radiation on whole-bone mechanical properties of mouse vertebrae from 20-week old mice, reported reductions in bone compressive monotonic strength and fatigue life at higher (above 17 000 Gy), but not at lower (50 and 1000 Gy) radiation doses [56]. Considering the very low radiation dosage (0.64 Gy per scan) used in our study, we do not expect reductions in bone mechanical properties. Nevertheless, the effects of long-term *in vivo* micro-CT imaging on bone mechanical properties should be addressed in future studies.

Interestingly, the heterogeneity in bone morphometry observed between imaging groups was not present for the FI measurements. At 40 weeks of age, the PolgA mice of the 5-scan group had significantly higher FI scores compared to those of the 2 scan group; however, as no differences were detected between the 11- and 2-scan groups, we expect that this difference was related to variation between animals rather than to the cumulative effects of radiation and anesthesia associated with *in vivo* micro-CT imaging. The imaging groups did not show any differences in body weights, which increased over time both in PolgA and WT mice. The absence of weight loss further supports the fact that the well-being of the animals was not negatively affected by the higher number of *in vivo* micro-CT scans in the first group, compared to groups 2,3 and 4, respectively. These results demonstrate how the parallel tracking of the FI as an addition to longitudinal *in vivo* micro-CT imaging not only allows to link age-related changes in bone morphometry to the development of frailty but also provides a useful tool to assess whether treatments/interventions have biasing effects on the overall health status of the animals. By maximizing the data obtained from individual animals, the total number of animals can be reduced. Based on the results of this study, we therefore recommend that in addition to acquiring baseline measurements of body weight and of the bone micro-architecture, baseline FI measurements should be included for studies in aging mice. Stratifying the animals according to these initial measurements could also be highly valuable to reduce the variability between study groups.

In conclusion, the combination of the longitudinal assessments of the FI and time-lapsed *in vivo* micro-CT imaging allowed for the detection of hallmarks of osteopenia and aging across multiple systems. Although the long-term monitoring approach can potentially lead to small but significant changes in bone morphometric parameters, the comparison between genotypes was not impaired. Moreover, more frequent *in vivo* micro-CT imaging did not negatively affect the multi-system hallmarks of aging such as body weight and frailty index. In line with the goal of “Reduction”, the second principle of the 3R’s, long-term *in vivo* micro-CT imaging allows to reduce the number of animals required for experiments while maintaining sufficient statistical power to reach a valid conclusion and thus, provides a powerful tool for usage in aging studies.

## Acknowledgements

The authors gratefully acknowledge valuable inputs from Dr. Ilaria Bellantuono concerning the establishment of the frailty index at the institute for biomechanics.

## Supporting information

**Table S1. The effects of imaging session number (group), genotype and the interaction effect (genotype*group) on bone morphometric parameters and frailty index (FI) at 40 weeks of age were compared via two-way ANOVA analysis.** The p-values and effect sizes (*f*) of the main and interaction effects, respectively are listed below (significance level α =0.05).

**Table S2. P-values, effect sizes (*f*) and achieved power obtained by cross-sectional (one-way ANOVA) and longitudinal analysis (paired t-test)**. (Significance level α =0.05).

## Notes

### Competing Interest Statement

The authors have declared no competing interest.

### Summary of Updates

The methods have been revised to provide more details. Furthermore the introduction and discussion have been revised.

