## Supplementary information for "Effects of long-term *in vivo* micro-CT imaging on hallmarks of osteopenia and frailty in aging mice"

Prof. Ralph Müller, PhD

**Supporting information**

**Table S1**

**Table S2**

**Table S1. The effects of imaging session number (group), genotype and the interaction effect (genotype*group) on bone morphometric parameters and frailty index (FI) at 40 weeks of age were compared via two-way ANOVA analysis.** The p-values and effect sizes (*f*) of the main and interaction effects, respectively are listed below (significance level α =0.05).

| two-way ANOVA | **Interaction** | | **Group** | | **Genotype** | |
| --- | --- | --- | --- | --- | --- | --- |
|  | p value | effect size *f* | p value | effect size *f* | p value | effect size *f* |
| **BV/TV** | 0.081 | 0.31 | <0.0001 | 0.74 | 0.001 | 0.50 |
| **Tb.Th** | 0.061 | 0.33 | 0.005 | 0.46 | <0.0001 | 0.57 |
| **Ct.Ar/Tt.Ar** | 0.200 | 0.25 | 0.004 | 0.48 | <0.0001 | 0.84 |
| **Ct.Th** | 0.279 | 0.22 | 0.049 | 0.34 | <0.0001 | 0.86 |
| **BFR** | 0.475 | 0.17 | 0.008 | 0.44 | 0.005 | 0.39 |
| **BRR** | 0.411 | 0.28 | <0.05 | 0.44 | <0.0001 | 0.59 |
| **MAR** | 0.403 | 0.18 | 0.065 | 0.32 | 0.029 | 0.30 |
| **MRR** | 0.641 | 0.13 | 0.002 | 0.50 | <0.0001 | 1.21 |
| **FI** | 0.143 | 0.27 | 0.054 | 0.33 | <0.0001 | 1.09 |

| **Cross-sectional analysis** | one-way ANOVA | **WT** | | | **PolgA** | | |
| --- | --- | --- | --- | --- | --- | --- | --- |
|  |  | p-value | effect size *f* | achieved power | p-value | effect size *f* | achieved power |
|  | **BV/TV** | 0.007 | 0.71 | 0.88 | <0.0001 | 0.96 | 0.99 |
|  | **Tb.Th** | 0.087 | 0.48 | 0.53 | 0.001 | 0.75 | 0.97 |
|  | **Ct.Ar/Tt.Ar** | 0.030 | 0.58 | 0.72 | 0.088 | 0.41 | 0.52 |
|  | **Ct.Th** | 0.125 | 0.43 | 0.46 | 0.146 | 0.36 | 0.42 |
|  | **BFR** | 0.285 | 0.33 | 0.26 | 0.020 | 0.54 | 0.66 |
|  | **BRR** | 0.151 | 0.41 | 0.42 | 0.023 | 0.53 | 0.64 |
|  | **MAR** | 0.687 | 0.18 | 0.11 | 0.054 | 0.45 | 0.61 |
|  | **MRR** | 0.033 | 0.57 | 0.70 | 0.044 | 0.47 | 0.53 |
| **Longitudinal analysis** | paired t-test | **WT** | | | **PolgA** | | |
|  |  | p-value | effect size *f* | achieved power | p-value | effect size *f* | achieved power |
|  | **BV/TV** | <0.0001 | 0.52 | 0.99 | 0.005 | 0.45 | 0.99 |
|  | **Tb.Th** | <0.0001 | 0.86 | 1.00 | 0.014 | 0.39 | 0.96 |
|  | **Ct.Ar/Tt.Ar** | <0.0001 | 0.57 | 1.00 | 0.011 | 0.29 | 0.80 |
|  | **Ct.Th** | <0.0001 | 0.91 | 1.00 | 0.002 | 0.36 | 0.92 |
|  | **BFR** | 0.050 | 0.59 | 1.00 | 0.094 | 0.32 | 0.85 |
|  | **BRR** | 0.091 | 0.43 | 0.93 | 0.314 | 0.22 | 0.56 |
|  | **MAR** | 0.698 | 0.11 | 0.15 | 0.988 | 0.01 | 0.05 |
|  | **MRR** | 0.019 | 0.52 | 0.99 | 0.610 | 0.10 | 0.15 |

**Table S2. P-values, effect sizes (*f*) and achieved power obtained by cross-sectional (one-way ANOVA) and longitudinal analysis (paired t-test)**. (significance level α =0.05).
